# Second-site capsule-locus mutations, not Ade efflux disruption, determine phage susceptibility across an *Acinetobacter baumannii* transposon mutant panel

**DOI:** 10.64898/2026.11.04.764408

**Authors:** Anita Tarasenko, Bhavya Papudeshi, Abbey L. K. Hutton, Maoge Zang, Brynley J. Davies, George Bouras, Vijini Mallawaarachchi, Bart A. Eijkelkamp, Robert A Edwards

## Abstract

Resistance-Nodulation-Division (RND) efflux pumps have been implicated in phage adsorption in several Gram-negative bacteria, suggesting that AdeABC, AdeFGH and AdeIJK, the three characterised RND efflux systems of multidrug-resistant *Acinetobacter baumannii,* could play a similar role. We tested this hypothesis using five phages (Equilibrator, Taro, AT01, BB02, and AC03) and a panel of 13 *A. baumannii* strains, including nine AB5075 transposon mutants affecting AdeABC, AdeFGH, or AdeIJK. Host-range testing, efficiency-of-plating assays, capsule-associated density measurements, antimicrobial susceptibility testing, and whole-genome sequencing were performed to assess the relationship between genotype and phage infection. Overall, the data did not support a role for the Ade components as receptors. Phages Equilibrator, Taro and AT01 formed plaques on every strain, and the restricted host ranges of BB02 and AC03 did not correspond to disruption of a particular efflux system. Sequencing instead identified second-site mutations in the KL25 capsule locus in six of the ten AB5075 strains. These included a 1,998-bp deletion affecting *wzc*, substitutions in *mnaA* and *mnaB*, an IS*Aba13* insertion in a putative glycosyltransferase and a subpopulation-associated frameshift in *wzx*. AC03 formed plaques on all six strains with capsule-locus changes but on none of the five strains without them. Capsule dependence differed among the other phages: relative to the parental AB5075, infection by these phages, Equilibrator, Taro and BB02, was restricted on the capsule-deficient *wzc*::T26 strain, whereas AT01 efficiency was only slightly reduced. Insertion-site mapping also distinguished a near-terminal *adeK* insertion that retained the parental antimicrobial phenotype from an early *adeK* insertion associated with loss of AdeIJK function. Together, these findings identify capsule-locus variation as the strongest correlation with phage susceptibility in this panel and demonstrate the importance of verifying mutant genomes before assigning phenotypes to their annotated insertions.

**Impact statement:** Transposon mutant collections are valuable for linking bacterial genes to phenotypes, but the recorded insertion does not necessarily describe every genetic change in a strain. We investigated whether the Ade efflux systems of *A. baumannii* contribute to phage susceptibility. Although the initial host-range patterns suggested differences among efflux mutants, whole-genome sequencing revealed independent capsule-locus mutations in six of ten AB5075 strains. Newly isolated phage AC03 formed plaques on every strain with a capsule-locus change and on none with wild-type capsule loci. This association explained differences between strains carrying separate insertions in the same *ade* genes. The other phages we isolated also differed in their dependence on capsule: some infected the capsule-deficient mutant poorly, whereas AT01 was comparatively unaffected. These results emphasise two considerations for phage studies. The capsule state of the isolation or test host can strongly influence which phages are recovered and how efficiently they plate. Whole-genome verification is equally important when interpreting phenotypes from ordered mutant libraries, particularly when surface structures are involved.

## 1. Introduction

Multidrug-resistant (MDR) *Acinetobacter baumannii* remains a major challenge in healthcare settings. The World Health Organisation identifies it as a critical-priority pathogen, and severe infections can have mortality rates above 40%^1–3^. It causes ventilator-associated pneumonia, bloodstream infections and surgical-site infections, while resistance to last-resort agents continues to narrow treatment options^4^.

Efflux pumps are membrane transport proteins that recognise compounds that enter the cell and expel them before they accumulate to inhibitory concentrations at intracellular targets. Because a single pump can accept structurally unrelated substrates, efflux is one of the few resistance mechanisms that acts against several antibiotic classes at once. In Gram-negative bacteria, the clinically dominant family is the Resistance-Nodulation-Division (RND) superfamily^5,6^. RND pumps are tripartite assemblies (Figure 1): an inner-membrane transporter captures substrate from the cytoplasm or the inner leaflet of the membrane and drives export using the proton-motive force; a periplasmic adaptor protein bridges the periplasmic space; and an outer-membrane channel delivers substrate directly to the cell exterior. Because this architecture bypasses the periplasm entirely, RND efflux, combined with the low permeability of the Gram-negative outer membrane, produces high intrinsic resistance. The outer-membrane channel may also serve as a phage receptor because it is large, abundant and exposed on the cell surface.

**Figure 1:**
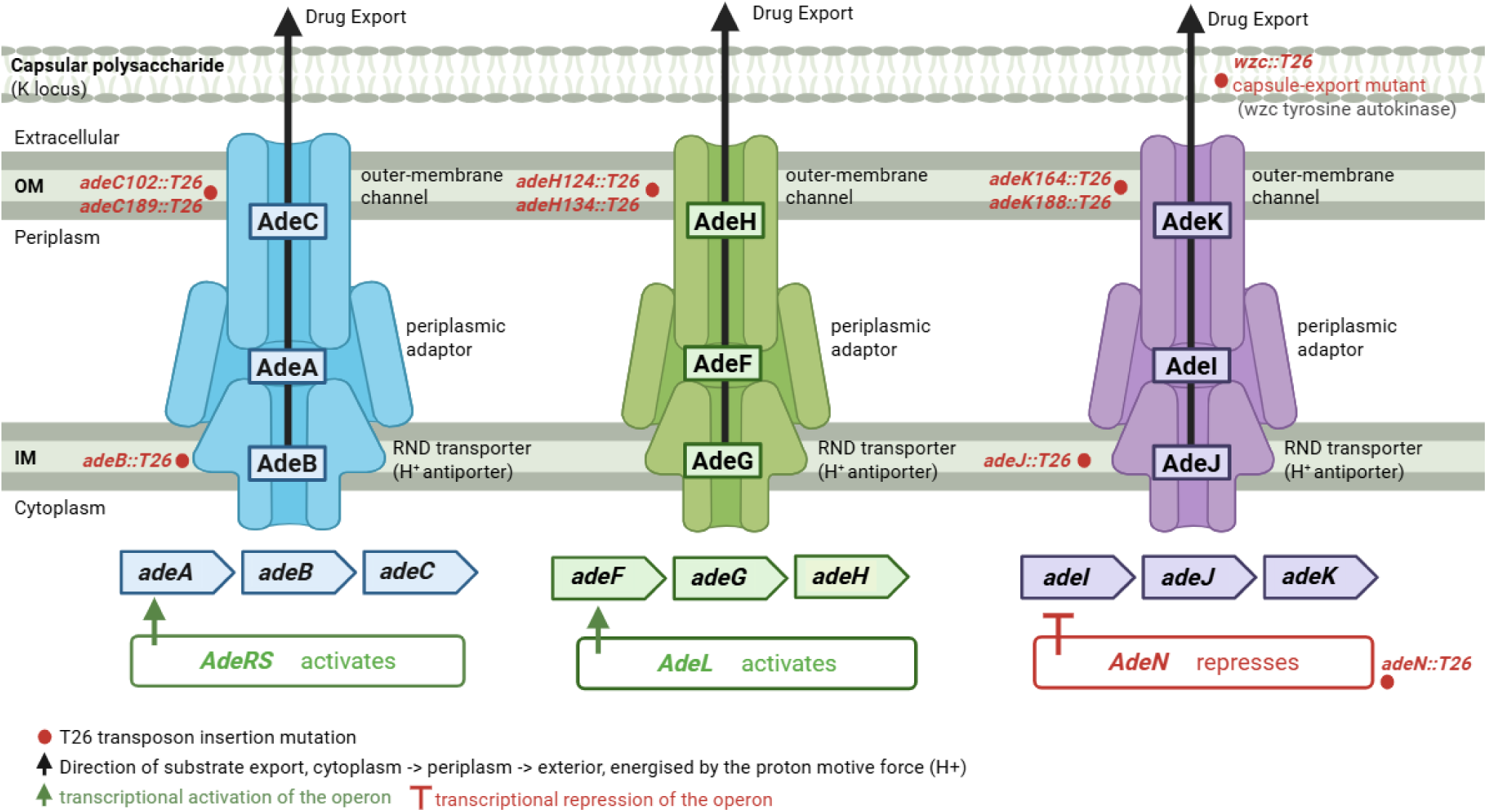
The three RND multidrug efflux systems of *A. baumannii*. Each system is a tripartite complex spanning the cell envelope, comprising an inner-membrane RND transporter energised by the proton-motive force, a periplasmic adaptor protein, and an outer-membrane channel through which substrates are delivered to the exterior. The AdeABC system is activated by the AdeRS two-component system encoded divergently upstream; the AdeFGH system is activated by the LysR-type regulator AdeL; and the AdeIJK system is constitutively expressed and repressed by the TetR-type regulator AdeN, encoded at a separate chromosomal locus. Exported substrate classes are listed for each system. Red markers indicate the components disrupted by T26 transposon insertion in the mutant panel used in this study, together with the capsule-export mutant *wzc*::T26. The capsular polysaccharide layer encoded by the K locus lies outside the outer membrane.

*A. baumannii* encodes three characterised RND systems, which differ in their components, their regulation and their substrate ranges (Figure 1). AdeABC comprises the AdeB transporter, the AdeA adaptor, and the AdeC channel, and is controlled by the AdeRS two-component system encoded divergently upstream; it exports aminoglycosides, tetracyclines (including tigecycline), fluoroquinolones, chloramphenicol, trimethoprim, and macrolides, and its overexpression is a principal driver of clinical tigecycline resistance^5,7^. AdeC is dispensable in several strains because AdeAB can recruit alternative outer-membrane channels. AdeFGH comprises the AdeG transporter, the AdeF adaptor, and the AdeH channel, and is activated by the LysR-type regulator AdeL; it is weakly expressed in wild-type cells and confers resistance to fluoroquinolones, chloramphenicol, trimethoprim, clindamycin, tigecycline, and sulfamethoxazole when de-repressed^5,6^. AdeIJK comprises the AdeJ transporter, the AdeI adaptor, and the AdeK channel; it is constitutively expressed and the principal contributor to intrinsic multidrug resistance. The TetR-type regulator AdeN, encoded at a separate chromosomal locus, represses it, and AdeIJK overexpression is toxic^5,6^.

Efflux systems may also influence phage infection, which could be exploited therapeutically, as phages that use efflux channels as receptors may impose trade-offs that reduce antibiotic resistance. In *Escherichia coli*, phage TLS uses TolC, an outer-membrane channel shared by AcrAB and other RND pumps, as its receptor^14^. In *Pseudomonas aeruginosa*, OMKO1 binds OprM, a channel used by MexAB-OprM and MexXY-OprM^15^. Changes that prevent OMKO1 infection can therefore impair efflux and increase antibiotic susceptibility^15^. In parallel with efflux, *A. baumannii* is surrounded by a capsular polysaccharide layer that serves as a major virulence factor and a means of immune evasion^8,9^. The highly variable K locus encodes capsule structure and abundance, and this diversity directly governs phage susceptibility, as phages initiate infection by binding surface-exposed receptors such as outer membrane proteins, lipopolysaccharide, or the capsular polysaccharide itself^10,11^. Dense capsule layers or capsule structures incompatible with a phage’s receptor-binding proteins physically impede adsorption and block access to underlying receptors; conversely, specific capsule types can themselves serve as high-affinity receptors^12,13^. In *A. baumannii*, genome-wide transposon screens have identified the BfmRS-dependent capsule-modulating pathways as a central driver of phage infection efficiency^12^. Concurrently, the BfmRS system also regulates efflux pathways^16–18^. These observations thus shed light on the regulatory interdependence of critical bacterial surface structures, such as between capsule and membrane efflux proteins, and their distinct roles in governing phage-bacteria interaction.

In this study, we asked whether the Ade systems of *A. baumannii* act as receptors for phage binding. If an Ade component is required for infection, disrupting it should prevent or markedly reduce plaque formation, whereas mutants affecting the same receptor should also show comparable patterns. Outer-membrane channels were of particular interest because they are exposed at the bacterial surface. We isolated five new *A. baumannii* phages and examined them across 13 *A. baumannii* strains: three wild-type strains, the capsule-deficient AB5075 *wzc*::T26 mutant, and nine AB5075 insertion mutants representing all three Ade systems^19^. Contrary to expectations, some paired mutants differed sharply in phage susceptibility profiles. Whole-genome sequencing instead identified second-site mutations in the KL25 capsule locus in six of the ten AB5075 derivatives. These capsule-locus changes, rather than the annotated ade insertions, matched AC03 susceptibility: the phage formed plaques on all six derivatives carrying these changes and on none of the four without them. Taken together, these findings link capsule-locus variation to phage susceptibility in this panel and highlight the importance of whole-genome verification before attributing mutant phenotypes to annotated transposon insertions.

## 2. Results

### 2.1 A defined insertion panel spanning all three Ade systems

We selected nine efflux-associated mutants to represent different parts of the Ade complexes, rather than treating each pump as a single unit (Figure 1; Table S1). The inner-membrane transporters were represented by *adeB*::T26 and *adeJ*::T26, and the channels by paired insertions in *adeC*, *adeH*, and *adeK*; *adeN*::T26 also disrupts the AdeIJK repressor. Each of the three genes, *adeC*, *adeH*, and *adeK,* contained two different insertions, allowing independent confirmation of the observed phenotypes. All nine strains were derived from AB5075. Together with the capsule-deficient *wzc*::T26 mutant, these are the ten AB5075 derivatives referred to throughout; we treat the AB5075 parent separately as the reference strain.

### 2.2 Five phages spanning two virion morphotypes

We isolated five novel phages from separate enrichments using different *A. baumannii* hosts: Equilibrator and Taro were isolated on ATCC 17978, AT01 was isolated on HUMC1, BB02 was isolated on AB5075, and AC03 was isolated on AB5075 *wzc*::T26 (Tables S1 and S2). All the phages produced clear, circular plaques without halos (Figure 2A; Table S2). Plaques were approximately 1 mm in diameter for Equilibrator and Taro, 3 mm for AT01 and 2 mm for BB02 and AC03.

**Figure 2:**
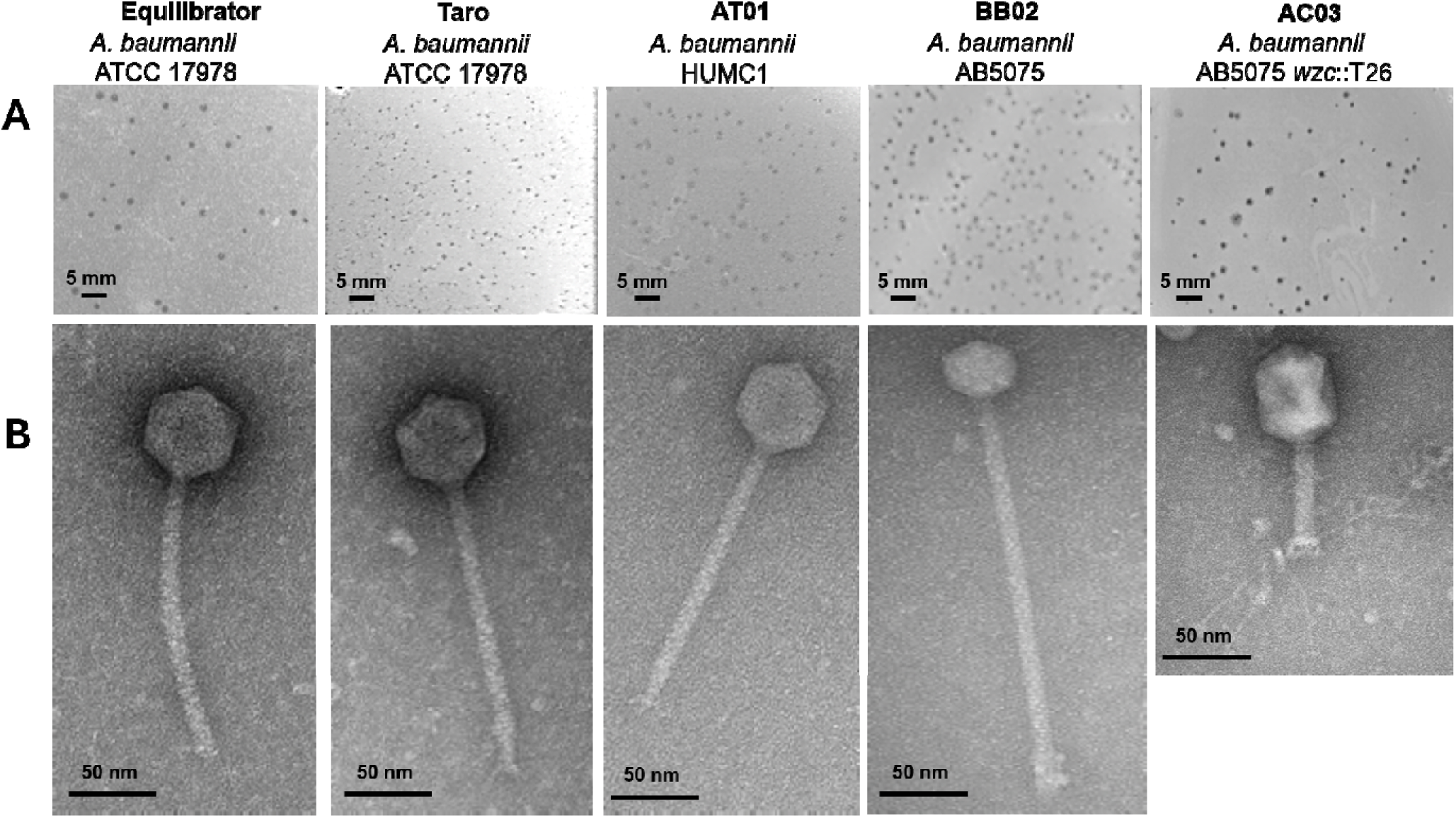
Plaque and virion morphology of five phages infecting *A. baumannii.* (A) Representative plaque morphology of Equilibrator and Taro on *A. baumannii* ATCC 17978, AT01 on HUMC1, BB02 on AB5075, and AC03 on the capsule-deficient AB5075 *wzc*::T26 mutant. Scale bars: 5 mm. (B) Representative transmission electron micrographs of negatively stained phage particles. Scale bars: 50 nm.

**Figure 3:**
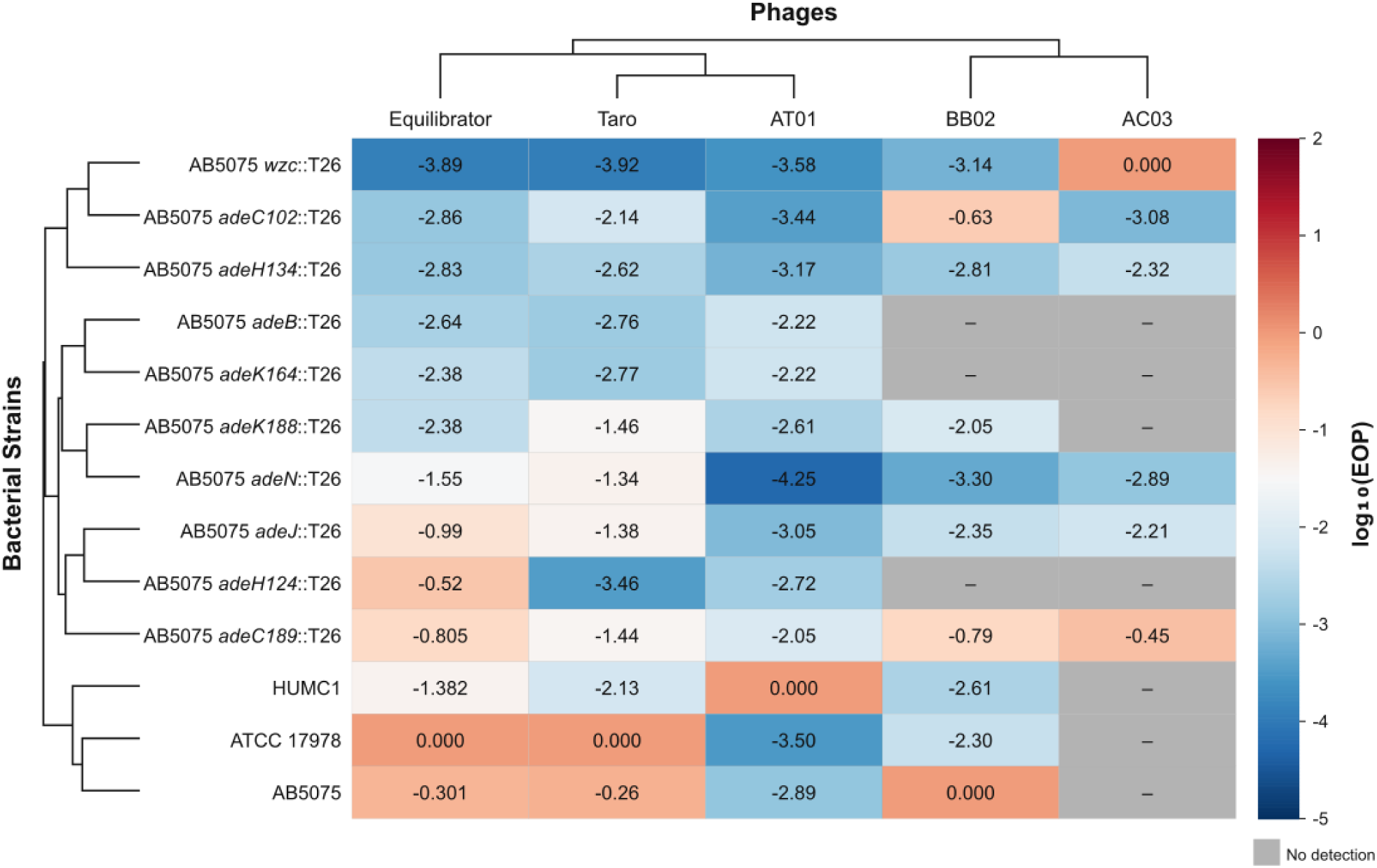
Hierarchically clustered log_10_(EOP) heatmap across *A. baumannii* strains. Rows represent bacterial genotypes and columns represent phages. Values are the mean log_10_(EOP) relative to each phage’s propagation host. Grey cells indicate no detectable plaques. The dendrograms on the left and above the heatmap represent hierarchical clustering of bacterial strains and phages, respectively, based on similarities in their log_10_(EOP) profiles. Bacterial strains were clustered by their susceptibility profiles across the five phages, and phages were clustered by their plating-efficiency profiles across the bacterial strains. Shorter dendrogram branches indicate greater similarity between profiles.

Transmission electron microscopy (TEM) showed long, flexible, non-contractile tails in Equilibrator, Taro, AT01, and BB02, consistent with siphovirus-like morphology, whereas AC03 displayed a shorter contractile tail, consistent with myovirus-like morphology (Figure 2B; Tables S2). Equilibrator, Taro, and AT01 had similar virion dimensions (∼66 nm isometric head, ∼200 nm tail) and genome sizes of 41,856, 43,021, and 43,022 bp, respectively. BB02 had a smaller head (∼49.5 nm), the longest tail in the panel (∼346.5 nm), and the largest genome (78,866 bp). AC03 had the largest capsid (92.4 nm), a ∼115.5 nm contractile tail and a 43,214 bp genome. Genome GC content ranged from 40.25% to 46% (Table S3). Sequence-, structure-, and synteny-guided annotation identified no integrase or repressor genes in any assembly, consistent with the clear-plaque phenotypes.

### 2.3 Host range does not follow Ade system identity

We first screened plaque formation across the 13 bacterial strains (Table 1). Equilibrator, Taro and AT01 formed plaques on every strain, including all nine *ade* insertion mutants. Thus, none of the disrupted Ade components was essential for these phages to form detectable plaques. BB02 plated on 10 of 13 strains, with no plaques detected on *adeB*::T26*, adeH124*::T26 or *adeK164*::T26. AC03 formed plaques on six strains but not on the three wild-type isolates or on *adeB*::T26, *adeH124*::T26, *adeK164*::T26 and *adeK188*::T26.

**Table 1:** Qualitative plaque formation and strain genotype across the panel. Note: + indicates plaque formation; - indicates no detectable plaques. Lineages refer to the three intergenic lineage-marker SNPs (LM1-LM3) that distinguish two sub-populations arising during library construction (Table S8). AC03 forms plaques on every strain with a capsule-locus change and on none without one. Full genotype nomenclature is given in Table S9.

| Strain | Ade component | Second-site capsule-locus change | Lineage | Equilibrator | Taro | AT01 | BB02 | AC03 |
|---|---|---|---|---|---|---|---|---|
| ATCC 17978 | Wild-type | n/a (different strain) | - | + | + | + | + | - |
| HUMC1 | Wild-type | n/a (different strain) | - | + | + | + | + | - |
| AB5075 | Wild-type (parental) | None detected | A | + | + | + | + | - |
| AB5075 *wzc*::T26 | Acapsular | 1,998bp deletion in *wzc* | A | + | + | + | + | + |
| AB5075 *adeC102*::T26 | AdeABC channel | *mnaA* Thr98Pro | B | + | + | + | + | + |
| AB5075 *adeC189*::T26 | AdeABC channel | *mnaB* Pro95Leu | A | + | + | + | + | + |
| AB5075 *adeK164*::T26 | AdeIJK channel | None detected | B | + | + | + | - | - |
| AB5075 *adeK188*::T26 | AdeIJK channel | None detected | A | + | + | + | + | - |
| AB5075 *adeH124*::T26 | AdeFGH channel | None detected | B | + | + | + | - | - |
| AB5075 *adeH134*::T26 | AdeFGH channel | *mnaB* Pro160Ser | B | + | + | + | + | + |
| AB5075 *adeB*::T26 | AdeABC transporter | None detected | B | + | + | + | - | - |
| AB5075 *adeJ*::T26 | AdeIJK transporter | ISAba13 in ABUW_3824 | B | + | + | + | + | + |
| AB5075 *adeN*::T26 | AdeIJK repressor | *wzx* frameshift (79% of population) | A | + | + | + | + | + |

These differences did not align with the identity of the disrupted Ade system. Within AdeABC, loss of the AdeB transporter abolished BB02 plaque formation. Yet, both *adeC* channel mutants, which should inactivate AdeABC, supported BB02 plaque formation at the highest efficiencies recorded for that phage outside its propagation host (EOP 2.37 × 10□¹ and 1.63 × 10□¹). Within AdeFGH, two insertions in the same gene gave opposite outcomes: *adeH124*::T26 blocked both restricted phages, while *adeH134*::T26 permitted both. Within AdeIJK, *adeK164*::T26 blocked both, while *adeJ*::T26, *adeK188*::T26 and *adeN*::T26 permitted BB02.

If an Ade component served as a receptor, we would predict it to be the surface-exposed outer-membrane channel, and its loss should be the most restrictive; instead, *adeC189*::T26 was the most permissive strain in the panel for three of the five phages. Therefore, these findings do not support the hypothesis that AdeC, AdeH or AdeK functions as a phage receptor.

The contrasting phenotypes of the two *adeH* mutants prompted sequencing the strains; before discussing those results, the difference in phage susceptibility was quantified.

### 2.4 Quantitative EOP across three biological replicates

We measured efficiency of plating (EOP) in three independent biological replicates and calculated relative to the phage’s propagation host (EOP = 1; Table S4; Table S5). Among interactions with detectable plaques, EOP ranged from 5.67 × 10 to 1, spanning approximately 4.25 orders of magnitude. Replicate log_10_(EOP) values generally differed by less than 0.2. No test strain plated more efficiently than its corresponding propagation host, although restriction varied considerably between strains.

A phage could therefore have a broad host range without plating efficiently on every host. Equilibrator formed plaques throughout the panel, but its EOP fell from 1 on ATCC 17978 to 1.29 × 10□□ on *wzc*::T26. AT01 likewise ranged from 1 on HUMC1 to 5.67 × 10□□ on *adeN*::T26. Taro plated relatively efficiently on parental AB5075 (5.55 × 10□¹), yet its EOP was lower on every mutant, ranging from 5.39 × 10□² on *adeN*::T26 to 1.19 × 10□□ on *wzc*::T26.

### 2.5 Genome sequencing reveals second-site capsule-locus mutations unrelated to the *ade* insertions

To characterise the genetic background of the mutant panel and the observed patterns of plaque formation and efficiency of plating, bacterial genomes were sequenced using Oxford Nanopore Technology PromethION and assembled *de novo.* This allowed verification of the expected insertion sites and identification of additional genomic variation that could contribute to differences in phage susceptibility (Tables S7–S13). The expected insertions were confirmed in every mutant. However, our genome sequencing also identified independent mutations in six of the ten AB5075 strains within the KL25 capsule biosynthesis and export locus, spanning *galU* (ABUW_3819) to *wzc/ptk* (ABUW_3833), that are not part of the original annotations and are unrelated to the accompanying *ade* insertions (Table 1; Table S9 and S10). These second-site changes affect four functional steps in capsule production. Comparative analysis of the KL25 locus across the AB5075 derivatives identified sequence variation affecting genes involved in capsule precursor synthesis, repeat-unit assembly and polysaccharide export (Table S10; Figure S1)

Three strains carry independent and distinct non-synonymous substitutions in the UDP-sugar precursor pathway: *adeC189*::T26 also contains MnaB Pro95Leu, *adeH134*::T26 also contains MnaB Pro160Ser, and *adeC102*::T26 also contains MnaA Thr98Pro. MnaA is a UDP-N-acetylglucosamine 2-epimerase, and MnaB is a UDP-N-acetylmannosamine dehydrogenase; together, they generate UDP-ManNAcA, a building block incorporated into the capsule repeat unit, and capsule-deficient *mnaA* and *mnaB* mutants have been reported previously^20^ (Table S10).

Separately, the *adeJ*::T26 strain carries an ISAba13 insertion at the 5’ end of ABUW_3824, a glycosyltransferase within the same locus. The *adeN*::T26 strain carries a 1-bp insertion in a homopolymeric run in *wzx* (ABUW_3827), extending the tract from eight to nine residues and inactivating the gene; read-level analysis showed 16% of reads retaining the parental eight-adenosine tract, 73% carrying nine and 6% carrying ten, with five remaining in 5% spanning other lengths or ambiguous at this position. Hence, approximately 79% of the population has an inactive *wzx* product (Table S10).

We identified four additional genomic features that are important for interpreting our results. First, three intergenic lineage-marker SNPs (LM1–LM3; Table S8) divide the strains into two sub-populations that evidently diverged during library construction: *adeB*::T26*, adeC102*::T26*, adeH124*::T26*, adeH134*::T26*, adeJ*::T26 and *adeK164*::T26 form lineage B. Meanwhile, the parent and the remaining derivatives retain the parental lineage A (Table 1 and Table S8). Second, *adeC102*::T26 is not a simple channel null: the T26 region is amplified to approximately three copies of the ABUW_1890–*adeC*′ segment, one intact *adeC* copy remains, and *adeAB* is duplicated. Third, the *adeK164*::T26 insertion occurs at the gene’s 3’ end and may not inactivate it. Finally, the AB5075 parent also differs from the AB5075-UW reference by a 7,510 bp insertion representing a second tandem copy of the Zot filamentous prophage, validated by long reads spanning the array (Table S7).

The *wzc*::T26 library yielded only 8x coverage and did not assemble, so this strain was assessed by mapping reads to the parental assembly rather than from a *de novo* assembly. Mapped reads support a 1,998 bp deletion in *wzc* (ABUW_3833, also annotated as the protein tyrosine kinase *ptk*), consistent with the intended capsule-deficient design. No other mutations were detected, but at this depth the read set does not support a confident claim that none are present.

### 2.6 Capsule-associated density phenotypes confirm capsule genotype

To test whether the capsule mutations we identified alter capsule expression levels, we assessed cell-surface phenotypes using a colloidal-silica density assay and TEM (Figure 4A–B, Table 2). The density assay provides an indirect, population-level measure of capsule-associated buoyant density rather than a direct quantification of capsule mass. Scored into four categories, the assay agreed with the capsule genotype in 10 of 11 AB5075 strains: all strains carrying a capsule-locus mutation scored none, low, or moderate. The *adeK188*::T26 strain scored high despite no detected capsule-locus change. Its genome was reconstructed at 22x overall sequencing coverage. Although no capsule-locus variant was detected, an undetected variant in this region cannot be ruled out.

**Figure 4:**
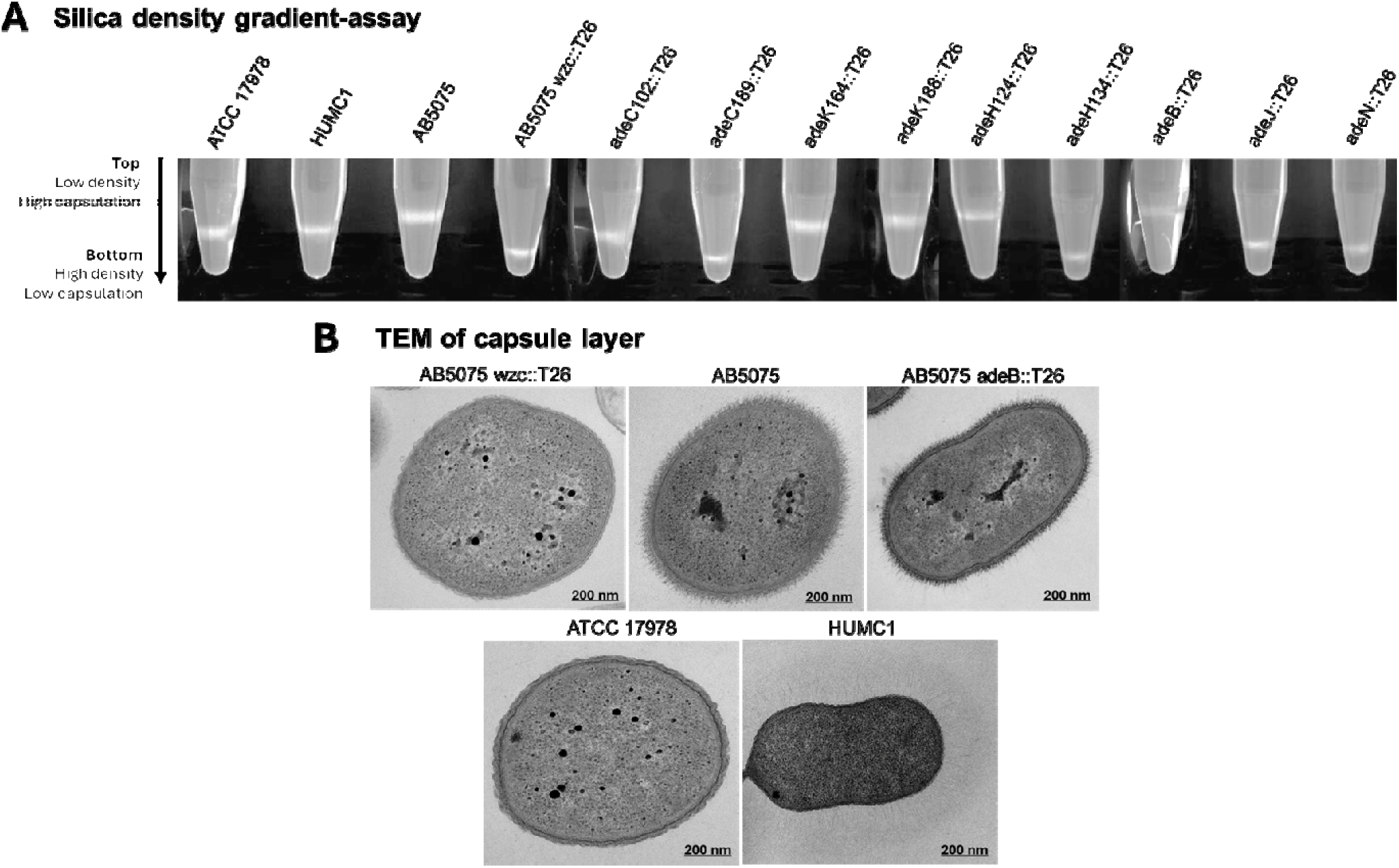
Capsule-associated phenotypes across *A. baumannii* strains. (A) Representative silica density gradient showing capsule-associated buoyant density. (B) TEM images of cell envelopes. Table 2 reports semi-quantitative capsule-associated buoyancy phenotypes based on gradient banding patterns in (A).

**Table 2:** Integrated capsule-associated phenotype, graded phage plaquing efficiency, and antimicrobial susceptibility across *A. baumannii* strains. Capsule score (None, Low, Moderate, High) reflects the semi-qualitative buoyancy phenotype shown in Figure 4A. Phage plaquing efficiency is graded as follows: +++ (EOP ≥ 0.5, including the propagation host, marked “ref”), ++ (0.1-<0.5), + (0.001-<0.1), ± (<0.001 but detectable), or - (no detectable plaques). MN, minocycline; TGC, tigecycline; AK, amikacin; CIP, ciprofloxacin. Full MIC data for all antibiotics tested are in Table S6.

| Strain | Capsule score | Equilibrator | Taro | AT01 | BB02 | AC03 | MN (µg/mL) | TGC (µg/mL) | AK (µg/mL) | CIP (µg/mL) |
|---|---|---|---|---|---|---|---|---|---|---|
| ATCC 17978 | Low | +++<br>(ref) | +++<br>(ref) | ± | + | - | 0.19 | 0.094 | 2 | 4 |
| HUMC1 | Moderate | + | + | +++<br>(ref) | + | - | 1.5 | 0.19 | >256 | >32 |
| AB5075 | High | +++ | +++ | + | +++<br>(ref) | - | 0.125 | 0.125 | 16 | 16 |
| AB5075 *wzc*::T26 | None | ± | ± | ± | ± | +++<br>(ref) | 0.125 | 0.125 | 16 | >32 |
| AB5075 *adeC102*::T26 | Moderate | + | + | ± | ++ | ± | 0.19 | 0.094 | 12 | >32 |
| AB5075 *adeC189*::T26 | Low | ++ | + | + | ++ | ++ | 0.125 | 0.094 | 24 | >32 |
| AB5075 *adeK164*::T26 | High | + | + | + | - | - | 0.125 | 0.125 | 12 | 16 |
| AB5075 *adeK188*::T26 | High | + | + | + | + | - | 0.047 | 0.047 | 4 | 3 |
| AB5075 *adeH124*::T26 | High | ++ | ± | + | - | - | 0.064 | 0.094 | 12 | >32 |
| AB5075 *adeH134*::T26 | Low | + | + | ± | + | + | 0.094 | 0.094 | 12 | 16 |
| AB5075 *adeB*::T26 | High | + | + | + | - | - | 0.047 | 0.023 | 2 | 3 |
| AB5075 *adeJ*::T26 | Low | ++ | + | ± | + | + | 0.047 | 0.064 | 8 | >32 |
| AB5075 *adeN*::T26 | Low | + | + | ± | ± | + | 0.038 | 0.19 | 12 | >32 |

Although the AB5075 transposon mutants were selected based on annotated insertions in Ade efflux-associated genes, whole-genome sequencing identified additional mutations in the KL25 capsule locus. These second-site changes provided an alternative explanation for the differences in phage susceptibility observed between strains carrying insertions in the same Ade component. Notably, AC03 formed plaques on all six strains with capsule-locus changes but on none without a detected change in this region. These findings suggest that AC03 susceptibility within the mutant panel was more closely associated with capsule-locus variation than with disruption of the Ade efflux system.

### 2.7 Capsule genotype predicts AC03 susceptibility

The strongest evidence that capsule rather than efflux governs susceptibility comes from AC03 (Table 3). Among the AB5075 derivatives, AC03 formed plaques on all six strains with a capsule-locus mutation and on none of the four without one. Capsule genotype therefore determines whether AC03 can infect at all, while the specific lesion modulates how efficiently it does so; efficiency among permissive strains ranged from 3.56 × 10□¹ on *adeC189*::T26 to 8.37 × 10□□ on *adeC102*::T26.

**Table 3:**
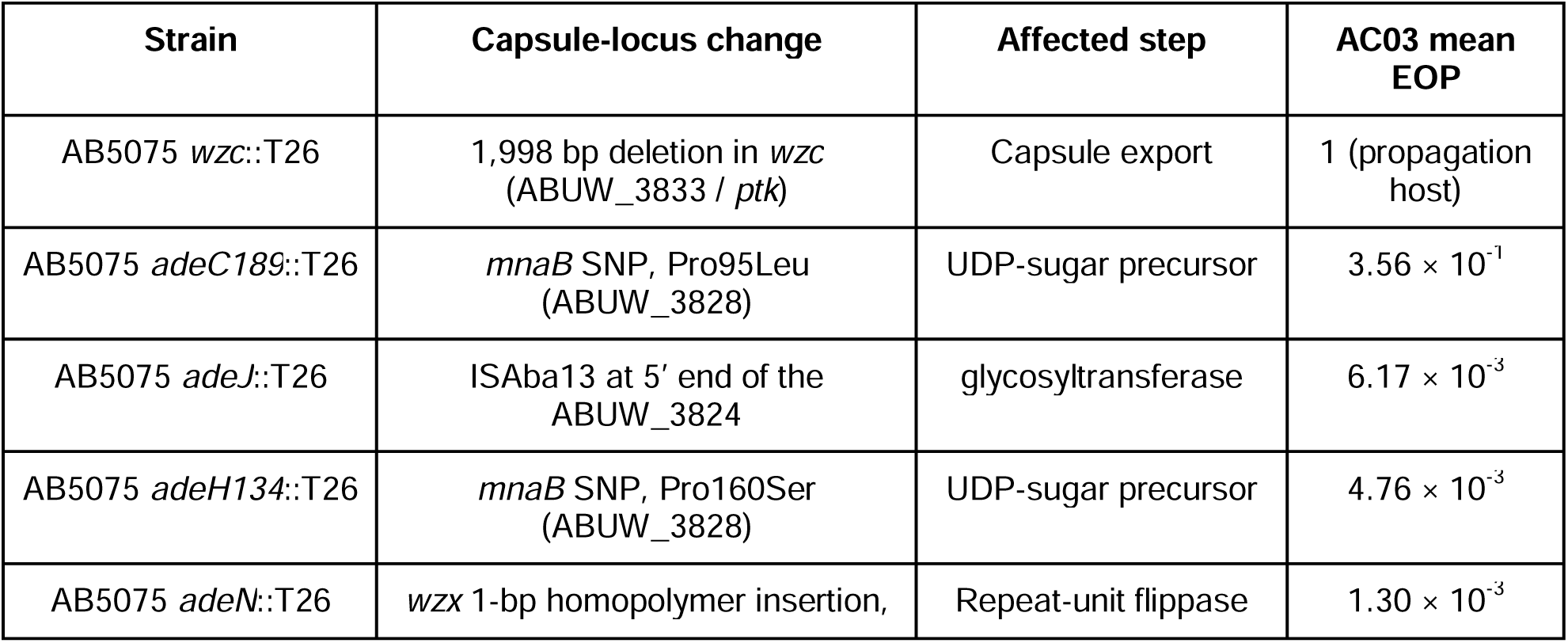
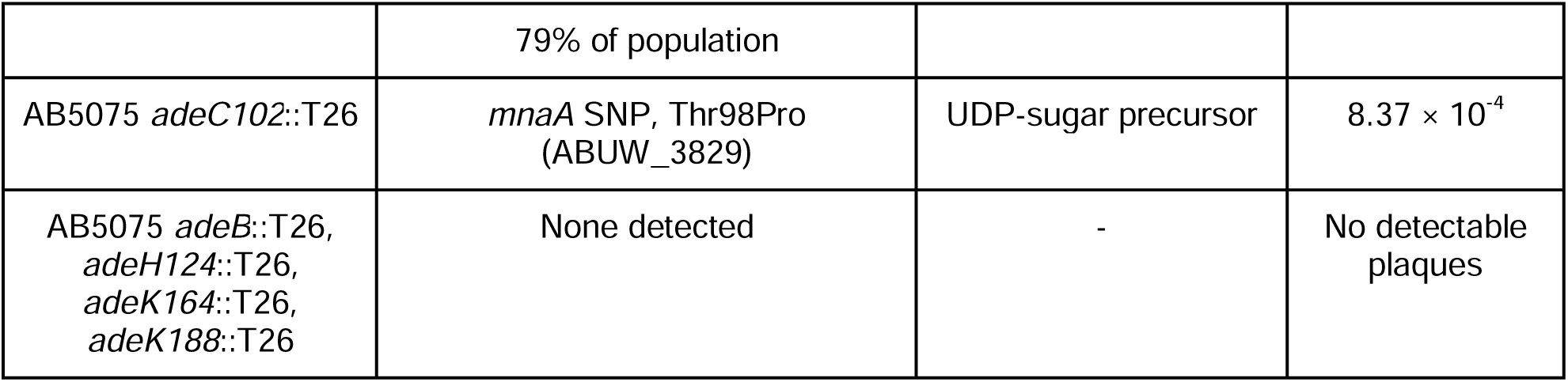
Capsule-locus genotype and AC03 EOP across the AB5075 derivatives.

The pattern did not reproduce when strains were grouped by the affected efflux system. The six permissive strains include insertions in all three Ade systems and in transporter, channel, and regulatory components. Although the panel did not include insertions in all nine structural genes across the three tripartite pumps, it sampled distinct components of each system: the AdeB and AdeJ transporters, the AdeC, AdeH and AdeK outer-membrane channels, and the AdeN regulators. The six non-permissive strains, by contrast, span multiple efflux systems and component types, rather than clustering by a single pump or component class. Conversely, the pairs of insertions in the same gene that appeared contradictory in Section 2.3 are immediately resolved: *adeC189*::T26 and *adeC102*::T26 both carry Mna substitutions, and both permit AC03. In contrast, *adeH134*::T26 carries MnaB Pro160Ser and permits AC03, but *adeH124*::T26 carries no capsule-locus change and does not. The phenotype tracks the second-site capsule mutation, not the *ade* insertion under which the strain is catalogued. AC03 fails to propagate on AB5075, HUMC1 and ATCC 17978, which carry different K-locus types, consistent with the same constraint operating across capsule types.

### 2.8 Capsule facilitates Equilibrator, Taro and BB02 infection; AT01 is largely indifferent

Three phages showed the opposite dependence to AC03. The isogenic comparison between AB5075 and its capsule-deficient strain *wzc*::T26 isolates the effect from background differences. Equilibrator fell from 5.00 × 10□¹ to 1.29 × 10□□ (3,876-fold), Taro from 5.55 × 10□¹ to 1.19 × 10□□ (4,664-fold), and BB02 from 1 to 7.18 × 10 (1,393-fold). Removing the capsule therefore restricts these three phages by three to four orders of magnitude, the signature expected if intact capsule facilitates adsorption or serves as the receptor, and the opposite of a steric-barrier model in which capsule loss should improve access to an underlying receptor.

AT01 behaved differently. Its EOP fell only from 1.30 × 10□³ to 2.61 × 10□□ across the same isogenic pair (a 5.0-fold change), and its EOP on *wzc*::T26 is within 1.2-fold of that on ATCC 17978 (3.14 × 10□□). AT01’s variation across the panel is therefore independent of the capsule state. Structure- and synteny-guided annotation of the AT01 genome also identified no recognised capsule depolymerase, although 34 out of 62 predicted protein-coding sequences (55%) remain functionally unassigned.

Overall, the phage collection thus shows three distinct relationships to the same surface structure: Equilibrator, Taro, and BB02 require a capsule for effective infection; AC03 infection requires an altered or absent capsule; and AT01 infection is largely indifferent to it.

### 2.9 Insertion position within *adeK* determines whether AdeIJK function is retained

Sequencing provided a possible explanation for the contrasting phenotypes of the two *adeK* insertion mutants, neither of which had a detected capsule-locus change. In *adeK164*::T26, the transposon insertion lies 18 bp (six codons) from the 3′ end of the gene, leaving a near-complete coding sequence intact and suggesting that AdeK function may be retained. This interpretation is supported by minimum inhibitory concentrations (MICs), the lowest antibiotic concentration that prevents visible bacterial growth, for minocycline, tigecycline, and amikacin that were comparable to those of the parental strain. In contrast, the insertion in *adeK188*::T26 occurs 51 codons into the 485-codon gene, about 10% into the coding sequence, and is predicted to disrupt function. The *adeK164*::T26 mutant also carries a 102 bp deletion removing one of four identical tRNA-Gln genes; three copies remain elsewhere in the genome, but the functional effect of this deletion was not assessed.

To test whether *adeK164*::T26 disrupts AdeK function, we measured the MIC of several antibiotics effective against *Acinetobacter* (Table 2). The MIC profiles matched these predictions. *adeK164*::T26 is indistinguishable from parental AB5075 across the AdeIJK substrate range, with all values within one twofold dilution of the parent: minocycline 0.125 µg mL□¹ (wild-type 0.125 µg mL□¹), tigecycline 0.125 µg mL□¹ (wild-type 0.125 µg mL□¹), ciprofloxacin 16 µg mL□¹ (wild-type 16 µg mL□¹) with the only difference, amikacin 12 µg mL□¹ (wild-type 16 µg mL□¹), falling within the reproducibility of gradient diffusion and well short of the coordinated multi-substrate reduction seen in the confirmed loss-of-function strains. *adeK188*::T26 showed coordinated reductions across the same agents: minocycline 0.047 µg mL□¹ (2.7-fold below wild-type), tigecycline 0.047 µg mL□¹ (2.7-fold) and amikacin 4 µg mL□¹(4-fold), phenocopying the *adeJ*::T26 transporter mutant of the same system (minocycline 0.047 µg mL□¹, tigecycline 0.064 µg mL□¹, amikacin 8 µg mL□¹) (Table 2; Table S6). We note that minocycline, tigecycline and amikacin are the substrates for which both known AdeIJK loss-of-function strains respond concordantly; tetracycline is unchanged in both *adeJ*::T26 and *adeK188*::T26 and therefore does not report AdeIJK function in this background, so the low tetracycline MIC recorded for *adeK164*::T26 (0.50 µg mL□¹ against a parental 4) is not evidence of AdeIJK loss and most likely reflects an unrelated effect.

Separate insertions in the same gene were therefore not phenotypically equivalent, consistent with the general principle that a transposon insertion’s consequence depends on its position within the target gene. Notably, *adeK164*::T26 retains a functional AdeIJK system and shows no capsule-locus change; AC03’s failure to propagate on this strain is consistent with its association with capsule loss or alteration, whereas BB02’s failure to propagate remains unexplained by the mutation identified in this panel.

## 3. Discussion

This study investigated whether *A. baumannii* phages use the outer-membrane components of AdeABC, AdeFGH or AdeIJK as receptors^5,6,21^. The susceptibility patterns did not support this explanation. Equilibrator, Taro and AT01 infected every *ade* insertion mutant, whereas the restricted host ranges of BB02 and AC03 did not follow the identity of the disrupted pump or channel. Independent insertions in the same gene also produced conflicting phage susceptibility profiles. These findings prompted examination of the bacterial genomes rather than attributing the differences directly to efflux function.

ONT sequencing revealed independent mutations in the KL25 capsule locus^22^ in six of the ten AB5075 derivatives that were unrelated to the identified *ade* insertions. These regions affect several stages of capsule production, including MnaA- and MnaB-dependent precursor synthesis, glycosyltransferase-mediated repeat-unit assembly, and Wzc- and Wzx-mediated export^23^. The study’s main contribution is linking these specific capsule-locus regions to distinct phage susceptibility patterns. Although the capsule is an established phage target, the effects of specific capsule-biosynthesis lesions and their relationship to outer-membrane proteins such as RND efflux components remain poorly resolved in *A. baumannii*. These findings provide a basis for investigating these interactions, but the data do not establish that Ade proteins serve as receptors or that capsule and efflux components interact directly during infection.

The cleanest association involved AC03, which was isolated using the capsule-deficient *wzc*::T26 strain. AC03 formed plaques on AB5075 derivatives with detected KL25 changes but not on those without them; consistent with altered capsules being important for infection. A host-range pattern that appeared inconsistent when the strains were considered solely as efflux mutants therefore became interpretable when their capsule genotype was taken into account. This result also highlights the importance of host capsule state during phage enrichment. Under the conditions tested, enrichment using only encapsulated strains would not have recovered AC03. Accordingly, phage collections assembled with encapsulated clinical isolates may undersample phages whose infection is obstructed by the capsule.

The other phages showed different relationships with the capsule state. In the isogenic AB5075 *wzc*::T26 strain comparison, Equilibrator and Taro exhibited approximately three orders of magnitude lower EOP on the capsule-deficient mutant, suggesting that intact capsule facilitates efficient infection^10,24^. AT01 showed only a 5-fold difference, indicating substantially less sensitivity to capsule loss under the conditions tested. These contrasting responses are consistent with the diverse relationship between *A. baumannii* phages and the capsule^10,12,24,25^. However, because disrupting *wzc* may alter the bacterial envelope beyond capsule loss^16^, these results do not establish capsule-dependent attachment, receptor identity, or the infection mechanisms of any of the newly isolated phages.

BB02 showed a more complex pattern. Its EOP on *wzc*::T26 was approximately 1,393-fold lower than on parental AB5075, consistent with capsule enhancing efficient infection. Nevertheless, BB02 also failed to form detectable plaques on *adeB*::T26, *adeH124*::T26 and *adeK164*::T26, none of which had a detected KL25 mutation. Conversely, *adeH134*::T26, which carried an independent mnaB Pro160Ser substitution, remained permissive, though with low EOP. These results suggest that capsule state alone does not explain BB02 susceptibility, but they do not establish a direct role for Ade proteins: *adeK164*::T26 appears to retain AdeIJK function, and the two *adeH* mutants had contrasting phenotypes. Resolving these differences will require independently constructed, isogenic mutants to distinguish the effect of capsule lesions, *ade* disruptions and other strain-specific features.

The frequency and diversity of capsule-loss mutations also raise questions about their origin. Two homopolymeric tracts occurred within the 1,224-bp *wzx* gene, separated by 398 bp. In the parental sequence, the 5′ tract contains eight adenosines and the 3’ tract contains seven. Expansion of the 5′ tract from eight to nine adenosines was detected in approximately 79% of reads from the lineage A *adeN* stock, whereas expansion of the 3′ tract from seven to eight adenosines predominated in the lineage B stock. Both expansions are predicted to cause frameshifts. Their occurrence in different tracts and stocks is consistent with separate slippage events, although their timing and the contribution of pre-existing population variation remain unresolved.

Other lesions included distinct point mutations in *mnaA* and *mnaB* and a novel IS*Aba13* insertion associated with loss of capsule. The diversity does not establish a site-specific mutation process or a mutator phenotype. The strains underwent several passages between their construction at the University of Washington in the early 2010s to their arrival at Flinders University, and any of these culturing steps could have enriched the capsule mutations. Similarly, Marshall *et al.*^20^ identified *mnaA*, *mnaB*, and *wzy* mutations, including spontaneous missense substitutions and IS*Aba13* insertions, in three spontaneous capsule mutants recovered from static AB5075 cultures. Capsule mutations appear to be frequently isolated in *A. baumannii*, suggesting they provide a growth advantage in common laboratory media. Marshall *et al.* also hypothesise that many of the original mutations accrued while “the original library [was] grown in deep well microtiter plates without aeration (Colin Manoil, personal communication)."

Insertion-site mapping provided a separate explanation for the functional differences between the two *adeK* mutants. The *adeK164*::T26 insertion lies six codons from the 3’ end and leaves AdeIJK functional, consistent with its parental MIC profile^26^. By contrast, *adeK188*::T26 interrupts the 485-codon gene a51 codons in and phenocopies the *adeJ*::T26 transporter mutant. Thus, transposon insertions do not necessarily represent a null allele; complicating library interpretation: the phenotype depends on what else has changed in the genome and where the insertion lies within the target gene (Table 2; Table S11). Moreover, AdeK can also interact with AdeAB independently of AdeC^27^.

Accordingly, the weaker antibiotic-resistance phenotype associated with *adeC* disruption than with *adeB* disruptions is compatible with AdeB retaining an alternative exit channel through AdeK, although the MIC data do not demonstrate this mechanism directly..

Together, these findings illustrate the need to verify the genotypes and functional status of strains obtained from ordered transposon libraries. Such libraries are often used on the assumption that each strain is different from its parent only at the catalogued insertion^19^. In this panel, however, second-site mutations are neither rare nor subtle, affecting six of ten strains carried additional capsule-locus mutations, including a 1,998 bp deletion, an IS insertion, and a frameshift, while three intergenic lineage markers indicated that the panel did not represent a single clonal lineage. Re-derivation and whole-genome sequencing are therefore important before assigning phenotypes to inferred genotypes, particularly for surface-associated phenotypes such as phage susceptibility, for which capsule variation is both common and consequential^28,29^. This study highlights, rather than replaces, the central biological finding that specific KL25 lesions are associated with distinct infection profiles.

Several limitations define priorities for further work. The capsule assay provides a semi-quantitative, population-level buoyancy measurement rather than a direct measure of capsule mass, although its results agreed with the capsule genotype in 10 of 11 strains. The capsule-deficient comparison also relied on the single *wzc*::T26 construct rather than independently generated capsule-null mutants. MIC measurements assessed cross-selection in pre-existing laboratory strains and were used to establish functional status, not to demonstrate an evolutionary trade-off under phage or antibiotic selection^30,31^. Biofilm formation was not tested, despite the involvement of both capsule^23^ and Ade systems^6,17,32^. Finally, these findings derive from a single strain background and its derivatives, and their generalisation to genetically diverse clinical isolates remains untested.

For phage therapy development, these emphasise the importance of characterising both the screening host and quantitative infection efficiency. Susceptibility patterns in efflux insertion mutants alone are insufficient to infer receptor usage^3,15^, particularly when capsule variation provides an alternative explanation^10,12^. Moreover, Equilibrator, Taro, and AT01 formed plaques on all 13 strains despite EOP differences of up to four orders of magnitude, demonstrating that qualitative host-range results can conceal substantial differences in infection efficiency. This is consistent with reports that single-dilution spot tests overestimate host range and virulence^33–35^. In this panel, capsule genotype and quantitative EOP were therefore more informative measures of phage susceptibility than the catalogued efflux-associated genotype alone.

## 4. Conclusion

The phage susceptibility patterns observed across this *A. baumannii* mutant panel were not explained by disruption of the AdeABC, AdeFGH or AdeIJK efflux systems. Instead, sequencing identified independent second-site mutations in the KL25 capsule locus in six AB5075 derivatives. These mutations matched AC03’s qualitative host range and explained several otherwise discordant results among strains carrying insertions in the same *ade* gene. The capsule’s influence varied across the phage collection. Equilibrator, Taro and BB02 plated substantially less efficiently on the capsule-deficient *wzc*::T26 strain, while AT01 was comparatively insensitive to the same change. Insertion position also mattered: the near-terminal *adeK164*::T26 insertion retained the parental antimicrobial phenotype, unlike the early *adeK188*::T26 insertion. Overall, our results show that capsule-associated genetic variation strongly correlates with phage susceptibility in this panel and underline the need to verify the full genomic background of transposon mutants before attributing a phenotype to a single annotated insertion.

## 5. Methods

### 5.1 Bacterial strain and culture conditions

Strains are listed in Supplementary Tables S1 and S8: *A. baumannii* ATCC 17978, HUMC1, AB5075, the capsule-deficient AB5075 *wzc*::T26 mutant, and nine efflux-associated T26 insertion mutants (Manoil Laboratory library)^19^. Glycerol stocks (-80 °C) were revived on LB agar (37 °C for 24 h) and grown in LB broth (37 °C, 180 rpm, 24 h) for liquid culture.

### 5.2 Phage isolation and purification

Wastewater was collected from the Glenelg Wastewater Treatment Facility (South Australia; 34°57′49″S, 138°30′38″E), clarified by centrifugation (4,500 *× g*, 10 min), and filtered through a 0.22 μm Sterivex filter (Millipore). Phages Equilibrator and Taro were isolated from ATCC 17978, AT01 from HUMC1, BB02 from AB5075, and AC03 on AB5075 *wzc*::T26, using double-agar overlay enrichment and screening^35^. Plaques were purified through three rounds of re-plating on the propagation host.

### 5.3 Host range and efficiency of plating (EOP)

Host range across the 13-strain panel was assessed by qualitative plaque formation; diffuse clearing without countable plaques was not scored as productive infection. For EOP, tenfold serial dilutions in SM buffer (50 mM Tris-HCl, 100 mM NaCl, 8 mM MgSO, pH 7.5) were plated with mid-log culture in molten LB top agar (37 °C, 18-24 h). EOP was calculated as titre on the test strain divided by titre on each phage’s respective propagation host (Equilibrator/Taro: ATCC 17978; AT01: HUMC1; BB02: AB5075; and AC03: AB5075 *wzc*::T26). Non-detected combinations were recorded as missing rather than assigned a pseudocount. Assays used three independent biological replicates (separate cultures, separate days) in technical triplicate; means across the three biological replicates are reported throughout, and individual replicate values are given in Table S4.

### 5.4 Colloidal silica density assay

Capsule-associated buoyant density was assessed by mixing normalised bacterial culture with LUDOX HS-40 colloidal silica, then centrifuging and recording band position as a qualitative phenotype scored into four categories (none, low, moderate, high). As buoyancy can reflect multiple cellular properties, this assay was treated as an indirect, population-level proxy for capsulation rather than a direct measure of capsule abundance. AB5075 *wzc*::T26 served as the capsule-deficient control.

### 5.5 Antimicrobial susceptibility testing

We determined MICs using gradient diffusion strips on Mueller-Hinton agar (0.5 McFarland, 37 °C, 18-24 h); raw observed values are reported, with the organism-specific breakpoint source stated for each antimicrobial. *Coli*stin gradient-diffusion results were retained only for descriptive purposes, as broth microdilution is the reference method for colistin susceptibility in *A. baumannii*^36^. For functional classification of insertion mutants, a mutant was considered loss-of-function when it showed coordinated reductions relative to the AB5075 parent across minocycline, tigecycline and amikacin, the three substrates for which both independently confirmed AdeIJK loss-of-function strains respond concordantly, and functional when its values across those substrates were indistinguishable from the parent.

### 5.6 Transmission electron microscopy (TEM)

Phage lysate (∼10^8^ PFU mL^-1^) was applied to plasma-cleaned, carbon-coated 200-mesh grids, stained with 1% uranyl acetate and lead citrate, and imaged on a Tecnai G2 Spirit TEM (FEI, 120 kV); morphology was measured from ≥10 virions/phage in ImageJ v1.53t. Bacterial samples were fixed (4% paraformaldehyde, 2.5% glutaraldehyde, 4 °C, 2 h), post-fixed (2% osmium tetroxide), dehydrated (70-100% ethanol), resin-embedded, and polymerised (70 °C, 48-72 h). Ultrathin sections (60-90 nm) were stained (4% uranyl acetate, lead citrate) and imaged at 100 kV.

### 5.7 Bacterial genome sequencing and assembly

Genomic DNA was extracted (NucleoSpin Tissue Mini Kit, Macherey-Nagel) and quantified (Qubit 1× dsDNA HS Assay, Thermo Fisher). Libraries were prepared with the Oxford Nanopore Rapid Barcoding Kit (SQK-RBK114-24) and sequenced on R10.4.1 flow cells (FLO-MIN114.24, MinION MK1D). Reads were basecalled with Slorado v0.6.0^37^ (super-accuracy R10.4.1), assembled with AutoCycler v0.8.0^38^, with per-strain read statistics, coverage and contig count reported in Tables S12 and S13. Coverage was calculated as read bases divided by the parental assembly total (4,074,487 bp, chromosome plus three plasmids) so that strains are directly comparable. Assemblies were annotated with Bakta v1.12.0^39^ plus structure-informed annotation via BaktFold v0.1.1^40^. Three caveats apply: *adeH124*::T26 and *adeK188*::T26 are single-assembler drafts; the seven contigs of *adeK188*::T26 include plasmid 1 broken into fragments and the BB02 phage genome, and the fifth contig of *adeK164*::T26 is likewise BB02; and *adeK164*::T26 was sequenced to 470× but assembled from a 35% subsample because the full read set was too large to assemble. The *wzc*::T26 library yielded 8× coverage and did not assemble.

### 5.8 Phage genome sequencing and assembly

Phage DNA was extracted (Norgen Phage DNA Isolation Kit) and quantified (Qubit 1× dsDNA HS Assay). Libraries were prepared (SQK-RBK114-24) and sequenced on R10.4.1 flow cells (FLO-MIN114.24, MinION MK1B). Reads were basecalled with Dorado v0.6.0 and quality-filtered at Q ≥ 10. Assembly and annotation used the Sphae v1.4.6 pipeline^41^, integrating Pharokka v1.9.1^42^, Phold v1.2.5^43^ and Phynteny-transformer v0.1.13^44^ (Table S3).

### 5.9 Insertion-site mapping and comparative genome analysis

Transposon insertion sites were determined from the assembled genomes and converted to predicted truncation points within each coding sequence. Our AB5075 wild-type was aligned end-to-end against the AB5075-UW reference (CP008706.1 and associated plasmids) to establish strain differences (Table S7). Variants in each strain were called relative to our wild-type rather than AB5075-UW, so second-site changes arising during library construction could be distinguished from pre-existing stock differences. Capsule-locus variants were identified within the KL25 locus spanning *galU* to *wzc* (ABUW_3819-3833). The tandem duplication of the Zot filamentous prophage in the wild-type strain was validated by long-read spanning: 19 reads span unique sequence through one full prophage copy into a second, 5 reads span the entire two-copy array, no reads support a one- or three-copy arrangement, and read depth is even across both copies (44.6×, 48.2× and 45.1× for the left flank, copy 1 and copy 2).

### 5.10 Competitive read mapping for phage DNA

Mapped bacterial read sets against a combined reference of the bacterial assemblies and five phage genomes to detect phage-derived DNA associated with each culture. Recorded genome coverage breadth, read counts, and phage-to-chromosome depth ratios for each strain. Host–phage junctions were searched for in split-read alignments to test for chromosomal integration, and split-read alignments between distinct positions within the phage reference were examined for evidence of circular or concatemeric molecules.

## 5.11 Data availability statement

Raw reads are deposited in NCBI SRA under BioProject PRJNA1353598. Reviewer link https://dataview.ncbi.nlm.nih.gov/object/PRJNA1353598?reviewer=5ak2p88ljeq6s2fmiiq8oof24b

## 6. Acknowledgements

The authors gratefully acknowledge the technical support and specialised expertise provided by Microscopy Australia at Adelaide Microscopy, Adelaide University, with support from NCRIS, the university, and state government funding. The authors thank SA Water (Glenelg) for providing wastewater samples. Flinders University’s DeepThought HPC, National Computational Merit Allocation Scheme grant (NCMAS-2026-28) from the Pawsey Supercomputing Research Centre, and the National Computational Infrastructure (NCI) supported the computational analyses.

## 8. Author Contributions

A.T.: Conceptualisation, Methodology, Investigation, Formal analysis, Data curation, Validation, Visualisation, Project administration, Writing: original draft, Writing: review & editing.

B.P.: Formal analysis, Bioinformatics, Data curation, Writing: review & editing.

A.H.: Investigation, Writing: review & editing.

M.Z.: Resources, Writing: review & editing.

B.D.: Methodology, Writing: review & editing.

G.B.: Bioinformatics, Data curation, Writing: review & editing.

V.M.: Investigation, Supervision, Funding acquisition, Writing: review & editing.

B.A.E.: Conceptualisation, Methodology, Resources, Supervision, Writing: review & editing.

R.A.E.: Conceptualisation, Methodology, Project administration, Formal analysis, Data curation, Bioinformatics, Supervision, Resources, Funding acquisition, Writing: review & editing.

## 10. Ethics Statement

No ethical approval was required for this study

