## Supplementary material for "Second-site capsule-locus mutations, not Ade efflux disruption, determine phage susceptibility across an *Acinetobacter baumannii* transposon mutant panel"

**Content**

1. Supplementary Figure (S1)
2. Supplementary Tables (S1 to S13)

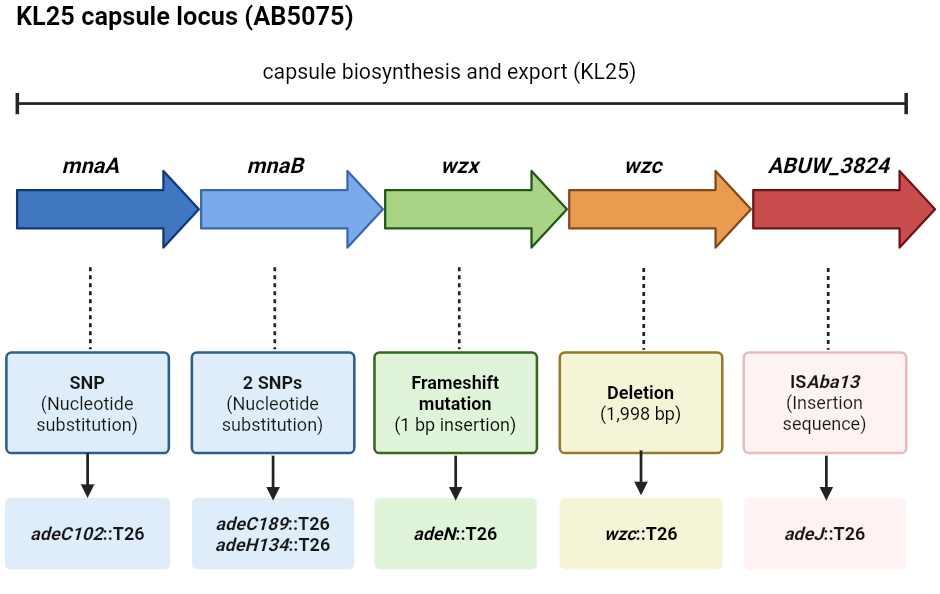

### **Figure S1: Schematic of the KL25 capsule locus in AB5075 and location of the second-site mutation identified in transposon mutants.** Arrows indicate predicted genes and their orientation. The mutation type and corresponding mutant strains are indicated beneath each affected gene. Table S10 provides gene locus tags, nucleotide substitutions, and genomic coordinates. Created in BioRender. Tarasenko, A. (2026) [https://BioRender.com/7ofrsfe](https://biorender.com/7ofrsfe).

### **Table S1: Genetic description of *Acinetobacter baumannii* strains and insertion mutants.**

| **Strain** | **Genetic strain/ description** | **Target gene** | **Locus tag** | **Genetic modification** |
| --- | --- | --- | --- | --- |
| ATCC 17978 | Reference strain; clinical isolate originally recovered from a case of meningitis | - | - | Wild type |
| HUMC1 | Clinical *A. baumannii* isolate | - | - | Wild type |
| AB5075 | Clinical isolate; parental background for T26 insertion mutant | - | - | Wild type |
| AB5075 *wzc*::T26 | Capsule-associated polysaccharide export mutant | wzc | ABUW_3833 | T26 transposon insertion |
| AB5075 *adeC102*::T26 | AdeABC-associated insertion mutant | adeC | ABUW_1976 | T26 transposon insertion (isolate 102) |
| AB5075 *adeC189*::T26 | AdeABC-associated insertion mutant | adeC | ABUW_1976 | T26 transposon insertion (isolate 189) |
| AB5075 *adeK164*::T26 | AdeIJK-associated insertion mutant | adeK | ABUW_0842 | T26 transposon insertion (isolate 164) |
| AB5075 *adeK188*::T26 | AdeIJK-associated insertion mutant | adeK | ABUW_0842 | T26 transposon insertion (isolate 188) |
| AB5075 *adeH124*::T26 | AdeFGH-associated insertion mutant | adeH | ABUW_1334 | T26 transposon insertion (isolate 124) |
| AB5075 *adeH134*::T26 | AdeFGH-associated insertion mutant | adeH | ABUW_1334 | T26 transposon insertion (isolate 134) |
| AB5075 *adeB*::T26 | AdeABC-associated insertion mutant | adeB | ABUW_1975 | T26 transposon insertion |
| AB5075 *adeJ*::T26 | AdeIJK-associated insertion mutant | adeJ | ABUW_0843 | T26 transposon insertion |
| AB5075 *adeN*::T26 | Insertion mutant targeting the transcriptional regulator adeN, associated with regulation of AdeIJK | adeN | ABUW_1731 | T26 transposon insertion |

### **Table S2: Plaque and virion characteristics of the five phages.**

| **Characteristics** | **Equilibrator** | **Taro** | **AT01** | **BB02** | **AC03** |
| --- | --- | --- | --- | --- | --- |
| **Isolation host** | ATCC 17978 | ATCC 17978 | HUMC1 | AB5075 | AB5075 wzc::T26 |
| **Plaque diameter** | ~1 mm (pinprick) | ~1 mm (pinprick) | ~ 3 mm | ~2 mm | ~2 mm |
| **Plaque shape** | Circular | Circular | Circular | Circular | Circular |
| **Plaque appearance** | Clear | Clear | Clear | Clear | Clear |
| **Halo formation** | No halo | No halo | No halo | No halo | No halo |
| **Virion morphology** | Long-tailed | Long-tailed | Long-tailed | Long-tailed | Short, contractile tail |
| **Head shape** | Isometric | Isometric | Isometric | Isometric | Isometric |
| **Head length (nm)** | ~66 | ~66 | ~66 | ~49.5 | ~115.5 |
| **Head width (nm)** | ~66 | ~66 | ~66 | ~49.5 | ~92.4 |
| **Tail length (nm)** | ~200 | ~200 | ~200 | ~346.5 | ~115.5 |
| **Tail width (nm)** | ~16.5 | ~16.5 | ~16.5 | ~13.2 | ~23.1 |
| **Total length (nm)** | ~264 | ~264 | ~264 | ~396 | ~231 |
| **Genome type** | dsDNA | dsDNA | dsDNA | dsDNA | dsDNA |

### **Table S3: Phage genome sequencing and assembly statistics.**

| **Phage** | **Equilibrator** | **Taro** | **AT01** | **BB02** | **AC03** |
| --- | --- | --- | --- | --- | --- |
| **Bioproject ID** | PRJNA1353598 | PRJNA1353598 | PRJNA1353598 | PRJNA1353598 | PRJNA1353598 |
| **Biosample ID** | SAMN53301081 | SAMN53301082 | SAMN53301083 | SAMN53301084 | SAMN53301085 |
| **SRA ID** | SRR36349990 | SRR36349989 | SRR36349988 | SRR36349987 | SRR36349986 |
| **Genome ID** | QB136406 | QB136410 | QB136407 | QB136409 | QB136408 |
| **Collection date** | 14 Aug 2023 | 14 Aug 2023 | 14 Aug 2023 | 14 Aug 2023 | 14 Aug 2023 |
| **Host** | ATCC 17978 | ATCC 17978 | HUMC1 | AB5075 | AB5075 wzc::T26 |
| **Number of reads (raw)** | 22,579 | 292,929 | 458,569 | 461,366 | 572 |
| **Total length (raw, bp)** | 122,533,676 | 571,292,826 | 513,371,554 | 866,865,825 | 1,215,612 |
| **Average read length (bp)** | 5,426.9 | 1,950.3 | 4,042 | 1,878.9 | 2,125.2 |
| **Genome size (bp)** | 41,856 | 43,021 | 43,022 | 78,866 | 43,214 |
| **Genome coverage (x)** | 2,927 | 13,280 | 11,932 | 10,991 | 28 |
| **Number of contigs** | 2 | 1 | 1 | 1 | 1 |
| **Completeness (%)** | 100.0 | 100.0 | 100.0 | 100.0 | 100.0 |
| **GC content (%)** | 46 | 45.61 | 43 | 40.25 | 46 |
| **Number of predicted CDS** | 62 | 62 | 62 | 123 | 154 |
| **CDS Coding Density** | 95.62 | 96 | 97.39 | 95.16 | 84.63 |
| **Read N50 (bp)** | 9,974 | 6,260 | 6,929 | 7,120 | 7,282 |
| **Read N75 (bp)** | 5,155 | 5,860 | 3499 | 3772 | 2322 |

### **Table S4: Individual biological replicate EOP values.**

| **Strain** | **Replicate** | **Equilibrator** | **Taro** | **AT01** | **BB02** | **AC03** |
| --- | --- | --- | --- | --- | --- | --- |
| ATCC 17978 | Rep 1 | 1 | 1 | 3.09 × 10⁻⁴ | 5.08 × 10⁻³ | – |
|  | Rep 2 | 1 | 1 | 3.56 × 10⁻⁴ | 4.92 × 10⁻³ | – |
|  | Rep 3 | 1 | 1 | 2.76 × 10⁻⁴ | 5.22 × 10⁻³ | – |
| HUMC1 | Rep 1 | 4.20 × 10⁻² | 7.22 × 10⁻³ | 1 | 2.38 × 10⁻³ | – |
|  | Rep 2 | 3.25 × 10⁻² | 8.79 × 10⁻³ | 1 | 2.88 × 10⁻³ | – |
|  | Rep 3 | 5.00 × 10⁻² | 6.15 × 10⁻³ | 1 | 2.09 × 10⁻³ | – |
| AB5075 | Rep 1 | 5.10 × 10⁻¹ | 5.56 × 10⁻¹ | 1.29 × 10⁻³ | 1 | – |
|  | Rep 2 | 4.58 × 10⁻¹ | 5.45 × 10⁻¹ | 1.52 × 10⁻³ | 1 | – |
|  | Rep 3 | 5.33 × 10⁻¹ | 5.64 × 10⁻¹ | 1.10 × 10⁻³ | 1 | – |
| AB5075 *wzc*::T26 | Rep 1 | 1.30 × 10⁻⁴ | 1.19 × 10⁻⁴ | 2.59 × 10⁻⁴ | 7.14 × 10⁻⁴ | 1 |
|  | Rep 2 | 1.25 × 10⁻⁴ | 1.21 × 10⁻⁴ | 2.95 × 10⁻⁴ | 8.14 × 10⁻⁴ | 1 |
|  | Rep 3 | 1.33 × 10⁻⁴ | 1.18 × 10⁻⁴ | 2.28 × 10⁻⁴ | 6.27 × 10⁻⁴ | 1 |
| AB5075 *adeC102*::T26 | Rep 1 | 1.40 × 10⁻³ | 7.22 × 10⁻³ | 3.60 × 10⁻⁴ | 2.38 × 10⁻¹ | 8.37 × 10⁻⁴ |
|  | Rep 2 | 1.33 × 10⁻³ | 7.27 × 10⁻³ | 4.09 × 10⁻⁴ | 2.20 × 10⁻¹ | 8.48 × 10⁻⁴ |
|  | Rep 3 | 1.44 × 10⁻³ | 7.18 × 10⁻³ | 3.24 × 10⁻⁴ | 2.54 × 10⁻¹ | 8.25 × 10⁻⁴ |
| AB5075 *adeC189*::T26 | Rep 1 | 1.89 × 10⁻¹ | 3.72 × 10⁻² | 8.92 × 10⁻³ | 1.62 × 10⁻¹ | 3.53 × 10⁻¹ |
|  | Rep 2 | 8.42 × 10⁻² | 3.97 × 10⁻² | 9.02 × 10⁻³ | 1.83 × 10⁻¹ | 3.15 × 10⁻¹ |
|  | Rep 3 | 1.97 × 10⁻¹ | 3.28 × 10⁻² | 9.03 × 10⁻³ | 1.45 × 10⁻¹ | 4.00 × 10⁻¹ |
| AB5075 *adeK164*::T26 | Rep 1 | 4.30 × 10⁻³ | 1.72 × 10⁻³ | 6.04 × 10⁻³ | – | – |
|  | Rep 2 | 3.83 × 10⁻³ | 1.76 × 10⁻³ | 6.74 × 10⁻³ | – | – |
|  | Rep 3 | 4.44 × 10⁻³ | 1.67 × 10⁻³ | 5.52 × 10⁻³ | – | – |
| AB5075 *adeK188*::T26 | Rep 1 | 4.20 × 10⁻³ | 3.44 × 10⁻² | 2.45 × 10⁻³ | 8.89 × 10⁻³ | – |
|  | Rep 2 | 3.25 × 10⁻³ | 3.94 × 10⁻² | 2.35 × 10⁻³ | 8.81 × 10⁻³ | – |
|  | Rep 3 | 5.00 × 10⁻³ | 3.00 × 10⁻² | 2.55 × 10⁻³ | 8.96 × 10⁻³ | – |
| AB5075 *adeH124*::T26 | Rep 1 | 3.10 × 10⁻¹ | 3.33 × 10⁻⁴ | 1.87 × 10⁻³ | – | – |
|  | Rep 2 | 2.83 × 10⁻¹ | 4.24 × 10⁻⁴ | 1.82 × 10⁻³ | – | – |
|  | Rep 3 | 3.11 × 10⁻¹ | 2.82 × 10⁻⁴ | 2.00 × 10⁻³ | – | – |
| AB5075 *adeH134*::T26 | Rep 1 | 1.50 × 10⁻³ | 2.42 × 10⁻³ | 6.76 × 10⁻⁴ | 1.56 × 10⁻³ | 4.74 × 10⁻³ |
|  | Rep 2 | 1.42 × 10⁻³ | 2.48 × 10⁻³ | 7.50 × 10⁻⁴ | 1.56 × 10⁻³ | 4.26 × 10⁻³ |
|  | Rep 3 | 1.56 × 10⁻³ | 2.33 × 10⁻³ | 6.14 × 10⁻⁴ | 1.55 × 10⁻³ | 5.28 × 10⁻³ |
| AB5075 *adeB*::T26 | Rep 1 | 2.30 × 10⁻³ | 1.72 × 10⁻³ | 6.04 × 10⁻³ | – | – |
|  | Rep 2 | 1.75 × 10⁻³ | 2.00 × 10⁻³ | 5.98 × 10⁻³ | – | – |
|  | Rep 3 | 2.78 × 10⁻³ | 1.51 × 10⁻³ | 6.07 × 10⁻³ | – | – |
| AB5075 *adeJ*::T26 | Rep 1 | 1.02 × 10⁻¹ | 4.28 × 10⁻² | 8.92 × 10⁻⁴ | 4.44 × 10⁻³ | 6.05 × 10⁻³ |
|  | Rep 2 | 8.08 × 10⁻² | 4.58 × 10⁻² | 8.94 × 10⁻⁴ | 5.25 × 10⁻³ | 5.22 × 10⁻³ |
|  | Rep 3 | 1.20 × 10⁻¹ | 3.79 × 10⁻² | 8.97 × 10⁻⁴ | 3.88 × 10⁻³ | 7.25 × 10⁻³ |
| AB5075 *adeN*::T26 | Rep 1 | 2.84 × 10⁻² | 5.33 × 10⁻² | 5.61 × 10⁻⁵ | 5.08 × 10⁻⁴ | 1.30 × 10⁻³ |
|  | Rep 2 | 2.30 × 10⁻² | 6.09 × 10⁻² | 6.29 × 10⁻⁵ | 4.92 × 10⁻⁴ | 1.30 × 10⁻³ |
|  | Rep 3 | 3.23 × 10⁻² | 4.74 × 10⁻² | 5.10 × 10⁻⁵ | 5.22 × 10⁻⁴ | 1.30 × 10⁻³ |

### **Table S5: Mean log_10_(EOP) of five phages across *A. baumannii* strains (n=3).**

| **Strain** | **Equilibrator** | **Taro** | **AT01** | **BB02** | **AC03** |
| --- | --- | --- | --- | --- | --- |
| ATCC 17978 | 0 | 0 | -3.50 | -2.29 | – |
| HUMC1 | -1.38 | -2.13 | 0 | -2.61 | – |
| AB5075 | -0.30 | -0.25 | -2.88 | 0 | – |
| AB5075 *wzc*::T26 | -3.89 | -3.92 | -3.58 | -3.14 | 0 |
| AB5075 *adeC102*::T26 | -2.86 | -2.14 | -3.44 | -0.63 | -3.08 |
| AB5075 *adeC189*::T26 | -0.80 | -1.44 | -2.05 | -0.79 | -0.45 |
| AB5075 *adeK164*::T26 | -2.38 | -2.76 | -2.21 | – | – |
| AB5075 *adeK188*::T26 | -2.38 | -1.46 | -2.61 | -2.05 | – |
| AB5075 *adeH124*::T26 | -0.52 | -3.46 | -2.72 | – | – |
| AB5075 *adeH134*::T26 | -2.83 | -2.62 | -3.17 | -2.81 | -2.32 |
| AB5075 *adeB*::T26 | -2.64 | -2.76 | -2.22 | – | – |
| AB5075 *adeJ*::T26 | -0.99 | -1.37 | -3.05 | -2.35 | -2.21 |
| AB5075 *adeN*::T26 | -1.55 | -1.27 | -4.25 | -3.29 | -2.89 |

### **Table S6: Antibiotic minimum inhibitory concentration (MIC; µg mL⁻¹) profiles.**

| **Strain** | **AMS** | **CAZ** | **FEP** | **MRP** | **CS** | **TE** | **MN** | **TGC** | **AK** | **CIP** | **C** |
| --- | --- | --- | --- | --- | --- | --- | --- | --- | --- | --- | --- |
| ATCC 17978 | 96 | >256 | >256 | 32 | 1.5 | 4 | 0.19 | 0.094 | 2 | 4 | >256 |
| HUMC1 | 16 | >256 | 32 | 48 | 1.5 | 12 | 1.5 | 0.19 | >256 | >32 | >256 |
| AB5075 | 64 | >256 | >256 | 16 | 1.5 | 4 | 0.125 | 0.125 | 16 | 16 | >256 |
| AB5075 *wzc*::T26 | 64 | >256 | >256 | 12 | 2 | 2 | 0.125 | 0.125 | 16 | >32 | >256 |
| AB5075 *adeC102*::T26 | 48 | >256 | >256 | 24 | 2 | 12 | 0.19 | 0.094 | 12 | >32 | >256 |
| AB5075 *adeC189*::T26 | 48 | >256 | >256 | 12 | 3 | 4 | 0.125 | 0.094 | 24 | >32 | >256 |
| AB5075 *adeK164*::T26 | 96 | >256 | >256 | 12 | 3 | 0.50 | 0.125 | 0.125 | 12 | 16 | >256 |
| AB5075 *adeK188*::T26 | 64 | >256 | >256 | 16 | 2 | 3 | 0.047 | 0.047 | 4 | 3 | 48 |
| AB5075 *adeH124*::T26 | 64 | >256 | >256 | 12 | 1.5 | 8 | 0.064 | 0.094 | 12 | >32 | >256 |
| AB5075 *adeH134*::T26 | 48 | >256 | >256 | 12 | 1.0 | 4 | 0.094 | 0.094 | 12 | 16 | >256 |
| AB5075 *adeB*::T26 | 48 | >256 | >256 | 16 | 1.5 | 4 | 0.047 | 0.023 | 2 | 3 | 96 |
| AB5075 *adeJ*::T26 | 64 | >256 | >256 | 16 | 1.5 | 4 | 0.047 | 0.064 | 8 | >32 | >256 |
| AB5075 *adeN*::T26 | 48 | >256 | 96 | 8 | 1.5 | 16 | 0.038 | 0.19 | 12 | >32 | >256 |

### *AMS = ampicillin-sulbactam; CAZ = ceftazidime; FEP = cefepime; MRP = meropenem; CS = colistin; TE = tetracycline; MN = minocycline; TGC = tigecycline; AK = amikacin; CIP = ciprofloxacin; C = chloramphenicol. Red = Resistant; Green = sensitive; Orange = intermediate.*

### **Table S7: Differences between the AB5075 parent used here and the AB5075-UW reference.**

| **Replicon** | **This study** | **AB5075-UW** | **Differences** |
| --- | --- | --- | --- |
| Chromosome | 3,980,185 bp | CP008706.1, 3,972,672 bp | 0 SNPs; 4 small insertions (9 bp); 6 small deletions (6 bp); one 7,510 bp insertion at CP008706.1:1,989,106 |
| Plasmid 1 | 83,604 bp | CP008707.1, 83,610 bp | 1 SNP; 6 small deletions (6 bp) |
| Plasmid 2 | 8,731 bp | CP008708.1, 8,731 bp | identical |
| Plasmid 3 | 1,967 bp | CP008709.1, 1,967 bp | identical |

### **Table S8: Lineage-marker SNPs distinguishing two sub-populations of the mutant panel.**

| **Label** | **Notation** | **Position in our assembly** | **Lineage A** | **Lineage B** | **Nearest gene** |
| --- | --- | --- | --- | --- | --- |
| LM1 | CP008706.1:g.2591354T>A | chromosome 2,598,774 | T | A | 52 bp upstream of ABUW_2598 |
| LM2 | CP008707.1:g.10287A>G | p1 10,287 | A | G | 157 bp upstream of ABUW_4020 |
| LM3 | CP008707.1:g.10292A>T | p1 10,292 | A | T | 162 bp upstream of ABUW_4020 |

### **Table S9: Genetic nomenclature of AB5075 derivatives based on whole-genome sequencing.**

| **Strain** | **Proposed genotype** | **Genetic background** |
| --- | --- | --- |
| AB5075 | *A. baumannii* AB5075 (parent stock; wild type at every locus below, but not identical to AB5075-UW) | A (parental lineage) |
| AB5075 *wzc*::T26 | AB5075 Δ*ptk*1004 | A (parental lineage) |
| AB5075 *adeB*::T26 | AB5075 *adeB1001::T26* | B (LM1–LM3 lineage) |
| AB5075 *adeC102*::T26 | AB5075 *adeC102::T26* ABUW_3829-1008 (T26 junction amplified, ~3 copies of the ABUW_1890–adeC′ segment; one intact adeC remains) | B (LM1–LM3 lineage) |
| AB5075 *adeC189*::T26 | AB5075 *adeC189::T26* ABUW_3828-1006 | A (parental lineage) |
| AB5075 *adeH124*::T26 | AB5075 ABUW_1334-124::T26 | B (LM1–LM3 lineage) |
| AB5075 *adeH134*::T26 | AB5075 ABUW_1334-134::T26 ABUW_3828-1007 | B (LM1–LM3 lineage) |
| AB5075 *adeJ*::T26 | AB5075 *adeJ1002::T26* ABUW_3824-1005::ISAba13 | B (LM1–LM3 lineage) |
| AB5075 *adeK164*::T26 | AB5075 *adeK164::T26* ΔtRNA-Gln | B (LM1–LM3 lineage) |
| AB5075 *adeK188*::T26 | AB5075 *adeK188::T26* | A (parental lineage) |
| AB5075 *adeN*::T26 | AB5075 ABUW_1731-1003::T26 wzx1009 | A (parental lineage) |

### **Table S10: Second-site capsule-locus (KL25) mutations and MnaA/MnaB alleles.**

| **Strain** | **Allele** | **Change** | **Position/locus tag** | **Affected step** |
| --- | --- | --- | --- | --- |
| AB5075 *wzc*::T26 | Δptk1004 | 1,998 bp deletion in *wzc* | ABUW_3833 | capsule export |
| AB5075 *adeC189*::T26 | ABUW_3828-1006 | MnaB Pro95Leu | g.3,902,719 G>A (JGHGPE_03805) | UDP-sugar precursor |
| AB5075 *adeH134*::T26 | ABUW_3828-1007 | MnaB Pro160Ser | g.3,902,525 G>A (JGHGPE_03805) | UDP-sugar precursor |
| AB5075 *adeC102*::T26 | ABUW_3829-1008 | MnaA Thr98Pro | g.3,903,875 T>G (JGHGPE_03806) | UDP-sugar precursor |
| AB5075 *adeJ*::T26 | ABUW_3824-1005::ISAba13 | IS insertion at 5' end | ABUW_3824 (JGHGPE_03801) | glycosyltransferase |
| AB5075 *adeN*::T26 | *wzx*1009 | 1-bp insertion in homopolymeric run (8→9) | *wzx* | repeat-unit flippase |
| AB5075, *adeB*::T26*,* *adeH124*::T26*, adeK164*::T26*, adeK188*::T26 | – | none detected | – | – |

### **Table S11: Functional interpretation of each strain from genome sequence.**

| **Strain** | **Interpretation** |
| --- | --- |
| AB5075 | Parental comparator; wild type at the loci under investigation. |
| AB5075 *wzc*::T26 | Probably capsule deficient. |
| AB5075 *adeB*::T26 | Likely non-functional AdeB. |
| AB5075 *adeC102*::T26 | Probably capsule deficient. Contains intact adeC and duplication of adeAB. |
| AB5075 *adeC189*::T26 | Probably capsule deficient. Likely non-functional AdeC. |
| AB5075 *adeH124*::T26 | Likely non-functional AdeH. |
| AB5075 *adeH134*::T26 | Probably capsule deficient and likely non-functional AdeH. |
| AB5075 *adeJ*::T26 | Probably capsule deficient and likely non-functional AdeJ. |
| AB5075 *adeK164*::T26 | Likely encodes a functional AdeK protein. |
| AB5075 *adeK188*::T26 | Likely non-functional AdeK. |
| AB5075 *adeN*::T26 | Likely non-functional AdeN. |

### **Table S12: Genome sequencing and assembly statistics for three wild-type and a capsule-deficient *A. baumannii* strains characterised in this study.**

| **Bacterial isolates** | **ATCC 17978** | **HUMC1** | **AB5075** | **AB5075 *wzc*::T26** |
| --- | --- | --- | --- | --- |
| **Bioproject ID** | PRJNA1353598 | PRJNA1353598 | PRJNA1353598 | PRJNA1353598 |
| **Biosample ID** | SAMN53301014 | SAMN53301015 | SAMN53301016 | SAMN53301017 |
| **SRA ID** | SRR36238872 | SRR36238871 | SRR36238870 | SRR36238869 |
| **Genome ID** | JCEPIW000000000 | JCEPIX000000000 | JCEPIY000000000 | JCEPIZ000000000 |
| **Collection date** | 12 Feb 2023 | 12 Feb 2023 | 12 Feb 2023 | 12 Feb 2023 |
| **Number of reads (raw)** | 52,808 | 17,022 | 60,964 | 11,586 |
| **Total length (raw, bp)** | 134,652,055 | 71,956,346 | 266,562,380 | 33,355,565 |
| **Average (bp)** | 2,549.8 | 4227.3 | 4,372 | 2,879 |
| **Genome size (bp)** | 4,809,160 | 4,144,257 | 4,074,487 | 3,978,607 |
| **Genome coverage (x)** | 32.9 | 17.4 | 65.5 | 8.3 |
| **Completeness (%)** | 100 | 100 | 100 | 100 |
| **GC content (%)** | 38.91 | 38.87 | 39.08 | 39.16 |
| **Number of predicted CDSs** | 3,634 | 3,846 | 3,749 | 184 |
| **tRNAs** | 73 | 72 | 74 | 1 |
| **tmRNAs** | 1 | 1 | 1 | 0 |
| **rRNAs** | 18 | 18 | 18 | 0 |
| **ncRNAs** | 34 | 30 | 32 | 0 |
| **Regulatory ncRNAs** | 7 | 10 | 7 | 0 |
| **CRISPR Arrays** | 0 | 0 | 3 | 0 |
| **oriCs/oriVs** | 1 | 2 | 1 | 0 |
| **oriTs** | 0 | 0 | 0 | 0 |
| **Pseudogenes** | 17 | 34 | 8 | 15 |

### **Table S13: Genome sequencing and assembly statistics for the nine efflux mutants of *A. baumannii* strains characterised in this study.**

| **Bacterial isolates** | **AB5075 *adeC102*::T26** | **AB5075 *adeC189*::T26** | **AB5075 *adeK164*::T26** | **AB5075 *adeK188*::T26** | **AB5075 *adeH124*::T26** | **AB5075 *adeH134*::T26** | **AB5075 *adeB*::T26** | **AB5075 *adeJ*::T26** | **AB5075 *adeN*::T26** |
| --- | --- | --- | --- | --- | --- | --- | --- | --- | --- |
| **Bioproject ID** | PRJNA1353598 | PRJNA1353598 | PRJNA1353598 | PRJNA1353598 | PRJNA1353598 | PRJNA1353598 | PRJNA1353598 | PRJNA1353598 | PRJNA1353598 |
| **Biosample ID** | SAMN53301018 | SAMN53301019 | SAMN53301020 | SAMN53301021 | SAMN53301022 | SAMN53301023 | SAMN53301024 | SAMN53301025 | SAMN53301026 |
| **SRA ID** | SRR36265879 | SRR36265878 | SRR36265877 | SRR36265876 | SRR36265875 | SRR36265724 | SRR36265723 | SRR36265722 | SRR36265721 |
| **Genome ID** | JCEPJA000000000 | JCEPJB000000000 | JCEPJC000000000 | JCEPJD000000000 | JCEPJE000000000 | JCEPJF000000000 | JCEPJG000000000 | JCEPJH000000000 | JCEPJI000000000 |
| **Collection date** | 12 Feb 2023 | 12 Feb 2023 | 12 Feb 2023 | 12 Feb 2023 | 12 Feb 2023 | 12 Feb 2023 | 12 Feb 2023 | 12 Feb 2023 | 12 Feb 2023 |
| **Number of reads (raw)** | 176,460 | 226,309 | 600,599 | 160,909 | 236,073 | 324,782 | 323,872 | 109,110 | 447,216 |
| **Total length (raw, bp)** | 435,115,704 | 938,997,157 | 1,913,828,008 | 91,287,483 | 238,398,178 | 526,748,440 | 751,168,126 | 367,973,525 | 752,420,288 |
| **Average read length (bp)** | 2,319 | 4,149 | 3,817 | 567 | 1010 | 1,622 | 2,319 | 3,372 | 1,682 |
| **Assembly (bp)** | 4,102,014 | 4,114,639 | 4,173,557 | 4,150,272 | 4,076,184 | 4,083,342 | 4,098,374 | 4,097,363 | 3,996,573 |
| **Genome coverage (x)** | 107 | 230 | 470 | 22 | 59 | 129 | 184 | 90 | 185 |
| **Read N50 (bp)** | 17,348 | 15,768 | 12,997 | 17,882 | 8,200 | 6,830 | 14,509 | 16,175 | 9,126 |
| **Number of contigs** | 4 | 4 | 4 | 7 | 4 | 4 | 4 | 4 | 4 |
| **Circular** | 4 | 4 | 4 | 3 | 2 | 4 | 4 | 4 | 4 |
| **Completeness (%)** | 100.0 | 100.0 | 100.0 | 100.0 | 100.0 | 100.0 | 100.0 | 100.0 | 100.0 |
| **GC content (%)** | 39 | 39 | 39.54 | 39.39 | 39.18 | 39.11 | 39.1 | 29.23 | 40 |
| **Number of predicted CDSs** | 3,931 | 30964 | 4,022 | 4,071 | 3,940 | 3,940 | 3,146 | 19 | 158 |
| **tRNAs** | 74 | 74 | 73 | 74 | 74 | 74 | 74 | 0 | 0 |
| **tmRNAs** | 1 | 1 | 1 | 1 | 1 | 1 | 1 | 0 | 0 |
| **rRNAs** | 18 | 18 | 18 | 18 | 18 | 18 | 18 | 0 | 0 |
| **ncRNAs** | 32 | 32 | 32 | 32 | 32 | 32 | 32 | 0 | 0 |
| **Regulatory ncRNAs** | 9 | 8 | 11 | 10 | 10 | 9 | 7 | 0 | 6 |
| **CRISPR Arrays** | 2 | 1 | 1 | 3 | 2 | 2 | 1 | 0 | 0 |
| **oriCs/oriVs** | 3 | 2 | 1 | 2 | 2 | 2 | 1 | 0 | 0 |
| **oriTs** | 0 | 0 | 2 | 0 | 0 | 0 | 0 | 1 | 0 |
| **Pseudogenes** | 4 | 4 | 3 | 9 | 0 | 6 | 3 | 0 | 2 |
